# Combining transcription factor binding affinities with open-chromatin data for accurate gene expression prediction

**DOI:** 10.1101/081935

**Authors:** Florian Schmidt, Nina Gasparoni, Gilles Gasparoni, Kathrin Gianmoena, Cristina Cadenas, Julia K. Polansky, Peter Ebert, Karl Nordström, Matthias Barann, Anupam Sinha, Sebastian Fröhler, Jieyi Xiong, Azim Dehghani Amirabad, Fatemeh Behjati Ardakani, Barbara Hutter, Gideon Zipprich, Bärbel Felder, Jürgen Eils, Benedikt Brors, Wei Chen, Jan G. Hengstler, Alf Hamann, Thomas Lengauer, Philip Rosenstiel, Jörn Walter, Marcel H. Schulz

## Abstract

The binding and contribution of transcription factors (TF) to cell specific gene expression is often deduced from open-chromatin measurements to avoid costly TF ChIP-seq assays. Thus, it is important to develop computational methods for accurate TF binding prediction in open-chromatin regions (OCRs). Here, we report a novel segmentation-based method, TEPIC, to predict TF binding by combining sets of OCRs with position weight matrices. TEPIC can be applied to various open-chromatin data, e.g. DNaseI-seq and NOMe-seq. Additionally, Histone-Marks (HMs) can be used to identify candidate TF binding sites. TEPIC computes TF affinities and uses open-chromatin/HM signal intensity as quantitative measures of TF binding strength. Using machine learning, we find low affinity binding sites to improve our ability to explain gene expression variability compared to the standard presence/absence classification of binding sites. Further, we show that both footprints and peaks capture essential TF binding events and lead to a good prediction performance. In our application, gene-based scores computed by TEPIC with one open-chromatin assay nearly reach the quality of several TF ChIP-seq datasets. Finally, these scores correctly predict known transcriptional regulators as illustrated by the application to novel DNaseI-seq and NOMe-seq data for primary human hepatocytes and CD4+ T-cells, respectively.

## 1 Introduction

Deciphering the system behind the complex regulation of gene expression in higher organisms is a challenging task in computational biology. A key aspect is to better understand the role of transcription factors (TFs), DNA binding proteins that regulate the transcriptional machinery in cells. TFs can activate and repress expression of genes that are located proximal or distal to their DNA binding site, by binding to promoters of genes or enhancers that are brought into close proximity via DNA looping [54]. TFs are known to have important roles in several diseases, e.g. a third of known human developmental disorders are related to deregulated TFs [64].

Several general approaches have been proposed to identify TFs acting as key players in gene regulation depending on the available data: Coexpression analysis combined with computational predictions of TF sequence binding can be used to identify key TFs [18]. Genome-wide TF binding data, as produced by TF ChIP-seq, is widely used to identify important TFs: ChIP-seq data was incorporated into coexpression analysis [43], was combined with transcriptome data [49, 66], used for the construction of Gene Regulatory Networks (GRNs) [8], and used together with Hi-C data [40].

Although ChIP-seq data delivers highly interpretable results, it is not well suited for high-throughput studies due to high costs and laborious procedures. Thus, current large epigenetic consortia such as Roadmap [39], Blueprint [1], and DEEP (http://www.deutsches-epigenom-programm.de/), do not generate TF-ChIP data. Instead, the generated epigenetic data is considered to predict TF binding, as it contains a wealth of information to simplify this task. Especially data on open-chromatin, as produced for example by DNaseI seq [34], ATAC-seq [6] or NOMe-seq [35], was shown to be well tailored for this purpose [47] and has become the standard for the analysis of tissue-specific TF binding in absence of TF-ChIP data. Using machine learning methods, these predictions can be used to identify TFs acting as key regulators [4, 46, 47].

In addition to open-chromatin data, also Histone Modification (HM) ChIP-seq data was used for the prediction of TF binding [4, 5, 13, 24, 51, 67]. In these studies, HMs were used either exclusively or along with open-chromatin data. It was shown that using DNaseI-seq data alone can lead to highly accurate TF binding predictions [13, 51], therefore we mainly focus on open-chromatin data in this article.

There are two general classes of methods to predict TF binding: site-centric methods [13, 32, 42, 51, 57, 69], and segmentation-based methods [3, 7, 24, 25, 27, 48, 50, 58].

Site-centric methods require the identification of putative TF binding sites (TFBS) using TF binding motifs represented with position weight matrices (pwms). According to the signal of the included epigenetic marks, the putative TFBSs are either classified as bound or unbound. There are various ways of incorporating epigenetic data in the prediction methods: In *Centipede*, not only open-chromatin information, but also histone modifications, genome conservation and the distance of a putative binding site to the closest TSS are considered using a hierarchical mixture model [51]. Another approach is taken in [13]. Here, an epigenetic prior is computed using the DNaseI-seq signal that is combined with a simple motif score. In the method *PIQ*, TFBS are predicted with Bayesian inference [57]. The supervised methods *MILLIPEDE* [42] and *BinDNase* [32] use a binned DNaseI-seq signal around candidate TFBS as features in a regression approach to predict truly active TFBS.

Segmentation-based approaches screen the DNaseI-seq signal for dips in DNaseI hypersensitive sites (DHS), so called footprints. These footprints are believed to be caused by TFs that are bound to DNA, thereby preventing the DNaseI enzyme from cutting [20]. Restricting the search space for active TFBS to footprints simplifies the prediction. There are methods based on sliding windows [48] as well as approaches based on hidden Markov models (HMM) [3, 25]. Using a binomial z-score, *DNase2TF*, interprets the depletion of DNaseI reads around putative footprints [58]. The method *Wellington*, uses a binomial test to identify footprints. A putative footprint is classified as a true footprint, if there are significantly fewer reads within it compared to its flanking region [50]. A subset of the footprint detection methods includes DNaseI bias correction [24], as the DNaseI cleavage bias was reported to affect the footprint calling [26, 36]. Unfortunately, footprinting methods have been applied mainly on DNaseI-seq data, but neither on ATAC-seq nor NOMe data. In addition, the possibility of segmenting based on peaks only, which are used for footprint detection, has not yet been systematically analysed and compared to the performance of footprint based segmentations. By considering only peaks, both ATAC-seq and NOMe data can be used easily, as the only required processing step is peak calling.

A drawback of all aforementioned approaches for TF binding prediction is the usage of hit-based motif screening algorithms, such as *Fimo* [21]. Hit-based methods use a threshold to decide whether a genomic site is considered to be a putative TFBS or not. Low affinity binding sites may be lost as they often do not pass the threshold. As it was shown that low-affinity binding is essential in biology [12, 59], this could negatively affect downstream analyses of TFBS. Here, we use a method called *TRAP*, that circumvents the drawback of hit-based methods by quantifying TF binding using a biophysically motivated model that produces binding affinity values for each TF [55]. TRAP affinity values for TFs have been shown to work well in the context of analysing co-regulated genes [56], ChIP-seq data and SNP analyses in TFBS [60], in gene expression learning [9] as well as in TF co-occurrence analysis using DHS regions [63].

We present a novel, generalizable, segmentation-based method, called *TEPIC*, to predict TF-binding using pwms, combined with a single open-chromatin assay. Using *TEPICs* predictions, we learn regression models to predict gene expression in several cell types using DNaseI-seq and NOMe-seq data. We show that using open-chromatin peaks performs favourably compared to footprints and that incorporating low-affinity binding enhances the quality of gene expression learning. In addition, we show that the signal of open-chromatin assays within peaks contains quantitative information that improves gene expression predictions further. Compared to previous work [46], *TEPIC* leads to better results and shows performance close to what is obtained using more expensive ChIP-seq data sets.

## 2 MATERIALS AND METHODS

### 2.1 Data

We apply *TEPIC* to data generated within the DEEP project, as well as to ENCODE data [15]. From DEEP, we use DNaseI-seq, and RNA-seq data for a HepG2 sample, DNaseI-seq and RNA-seq data for three biological replicates of primary human hepatocytes, as well as NOMe-seq and RNA-seq data for six CD4+ T-Cell samples, including different subtypes. From ENCODE, we downloaded DNaseI-seq data, gene expression data, H3K4me3 ChIP-seq data, H3K27ac ChIP-seq data, and TF-ChIP-seq files for K562, GM12878, and H1-hESC. For HepG2, we downloaded H3K4me3 ChIP-seq, H3K27ac ChIP-seq and TF-ChIP-seq files. In total, we obtained 33 TF ChIP-seq files for K562, 50 for GM12878, 50 for H1-hESCs, and 39 for HepG2. So, *TEPIC* is tested on both primary cells and cell lines. DEEP sample IDs and ENCODE accession numbers are listed in Supplementary Table 1. Within *TEPIC*, we tested two different sets of position weight matrices (pwms): Using Jaspar [44] and Uniprobe [30], we created a set of 439 pwms for usage within *TEPIC*. We downloaded the set of non redundant TFs for vertebrates from Jaspar (version of 26.10.2015) and vertebrate TF data from Uniprobe. In addition, we obtained the full set of human mononucleotide profiles from Hocomoco, version 10, which includes 641 pwms [38]. However, we found this collection of pwms to perform worse than the other one (see discussion, Supplementary Figure 2), thus we do not consider the Hocomoco pwms in the remainder of the manuscript. Another curated but only commercially available dataset of pwms from the TRANSFAC database was not considered in this work [45]. For further details on the origin and processing of the DEEP samples, we refer to Supplementary Section *Experimental Procedures*. The EGA accession number for the DEEP data used in this study is *EGAS00001002073*.

### 2.2 Data Preprocessing

Bedtools version 2.25.0 [53] has been used in several stages during preprocessing. Peak calling on DNaseI-seq data has been conducted with JAMM [31] using the suggested default parameters. JAMM takes bed files as input, which need to be generated from the original bam files. For downstream usage, we considered all peaks that passed the JAMM filtering step. NOMe peaks have been called using a HMM based approach (Nordström *et al.*, unpublished, available at *https://github.com/karl616/gNOMePeaks*).

For DEEP samples, BAM files of RNA-Seq reads were produced with TopHat 2.0.11 [61], with Bowtie 2.2.1 [41], and NCBI build 37.1 in `--library-type fr-firststrand` and `--b2-very-sensitive` setting. Gene expression has been quantified using Cufflinks version 2.0.2 [62], the hg19 reference genome and with the options `frag-bias-correct, multi-read-correct`, and `compatible-hits-norm` enabled.

Gene expression quantifications for K562, and GM12878, as well as HM peaks and TF ChIP peaks were used as obtained from ENCODE. We considered the mean gene expression in H1-hESC over four replicates, HM and TF-ChIP data were not modified.

### 2.3 TF annotation using TEPIC

We compute TF affinities within all identified open-chromatin regions/HM peaks using TRAP [55] on the pwm sets described above. The annotation is parallelised in R. TF affinities per gene are computed using python in four different ways: Summing up the TF affinities in all open-chromatin/HM peaks within [1] a 3000bp window around a genes TSS and [2] a 50000bp window around a genes TSS using exponential decay as introduced in [49]. In addition to the positional information of the peaks, we incorporate the signal abundance within a peak into the TF annotation by multiplying the average per-base read count within the peak (DNaseI-seq/HM) or the average methylation in the peak (NOMe-seq), by the TF affinities. We perform this for all peaks in [3] the 3000bp window and [4] the 50000bp window. In the remainder of the paper, we refer to [1] as the 3kb setup, to [2] as 50kb, to [3] as 3kb-S and to [4] as 50kb-S, where S is short for scaled. Formally, TF gene scores are computed as

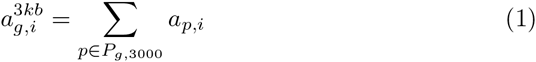

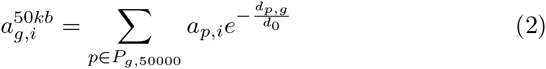

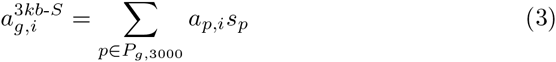

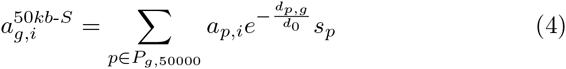

where [1]–[4] represent the previously described settings, *a*_*g,i*_ is the total affinity of TF *i* for gene *g, a*_*p,i*_ is the affinity of TF *i* in peak *p*, the set *P*_*g,x*_ contains all open-chromatin peaks in a window of size *x* around gene *g, d*_*p,g*_ is the distance from the centre of peak *p* to the TSS of gene *g, s*_*p*_ is the scaling factor used for peak *p*, and *d*_0_ is a constant fixed at 5000bp [49]. *TEPIC* is documented using a metadata xml file [16]. Each run automatically generates a meta analysis file containing all parameters used. The general workflow around *TEPIC* is shown in Figure 1.

**Figure 1.**
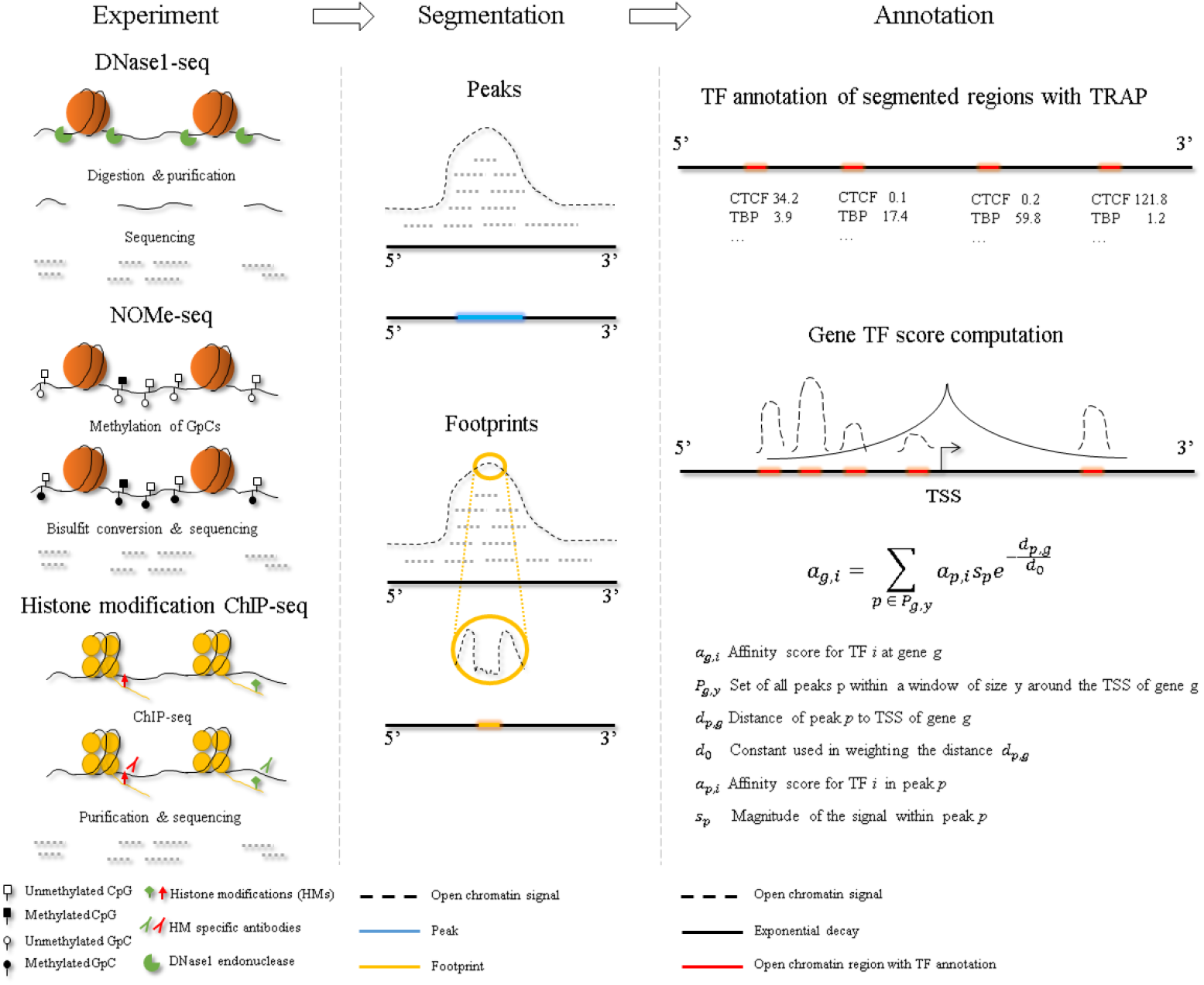
The general workflow of *TEPIC* is as follows: Data of an open-chromatin or Histone modification ChIP-seq experiment needs to be preprocessed to generate a genome segmentation, either by peak for footprint calling. Using the segmentation, TEPIC applies TRAP in all regions of interest, and computes TF gene scores using exponential decay to reweigh In addition, the magnitude of the open-chromatin signal is considered to reweigh TF scores in the segmented regions.

### 2.4 Elastic net regression to predict gene expression

We use the linear regression framework with elastic net penalty as implemented in the glmnet R-package [19] to predict gene expression from *TEPICs*, hit-based, and ChIP-seq TF binding predictions. As TFs are likely to be correlated, the elastic net is especially well suited for such a setting, because it resolves the correlation between features by distributing the feature weights among them [71]. This is achieved by combining two regularisation functions, the ridge penalty and the lasso penalty:

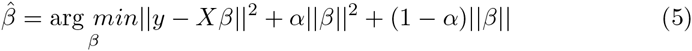

Here, *β* represents the feature coeffcient vector, 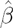 the estimated coefficients, *X* the feature matrix, and *y* the response vector. The ratio between lasso penalty and ridge penalty is controlled using the parameter *α*. Nested cross-validation is used to learn the models and to assess their performance.

In a ten-fold outer loop, we randomly select 80% of the data as training data and 20% as test data. On the training data, we perform a six-fold inner cross validation to learn model parameters. Within this step, we identify the optimal value for the parameter *α*, which is identified by a systematic search between 0.0 and 1.0 using a step-size of 0.01. The performance of the learned model is assessed on the hold-out test data. In the end, we report the average correlation *c*_*avg*_ on the test data sets over the ten-fold outer loop. Our learnig approach is further detailed in Supplementary Figure 1.

The data matrix *X*, containing TF gene scores, and the response vector *y*, containing gene expression values, are log-transformed, with a pseudo-count of 1, centered and normalised.

### 2.5 Competing TFBS prediction approaches

#### 2.5.1 Experimental using ChIP-seq

To compare our predictions to ChIP-seq data, we computed gene TF scores for all protein coding genes using ENCODE ChIP data and exponential decay as described in [49]. We considered all ChIP peaks within a window of 50000bp around the TSSs of genes.

#### 2.5.2 Segmentation with footprints

We obtained DNaseI-seq footprint predictions for HepG2, K562, GM12878, and H1-hESC generated with *HINTBC* [24]. As footprints can be shorter than the considered pwms, we extended the footprints to a total size of 24bps and 50bps, centered at the middle of the footprint. This data allows us to compare the peak-centric segmentation to the footprint-based segmentation. The extended footprint regions are annotated using all setups of *TEPIC*.

#### 2.5.3 Hit-based annotation methods

We applied the motif annotation tool *Fimo* [21] to open-chromatin peaks using the same set of pwms as we used for *TEPIC*. Thereby, we can assess the influence of the affinity-based binding prediction on gene expression learning. In addition, we run *Fimo* using a DNase prior [13], to compare *TEPIC* against a state of the art site-centric approach. This comparison is also motivated by the fact that this method has been used in a previous study on gene expression learning with TF binding predictions [46]. Transcription factor scores are computed in 3000bp and 50000bp windows around the transcription start sites (TSSs) of all protein coding genes. Gene TF scores are then calculated as described above for cases [1] and [2] using the log-ratio scores introduced in [13]. Note that we thus do not log-transform the Fimo TF scores in the elastic net model, comparable to [46]. We use the standard parameters of *Fimo* in all experiments, except for the `max-stored-scores` options which we set to 200000 instead of the default value 100000.

### 2.6 Evaluation using TF-ChIP-seq data

As it was noted previously, there is no standard procedure to compare TF binding predictions to ChIP-seq data [42]. Here, the gold standard dataset is constructed in a “peak-centric” manner: All *x* ChIP-seq peaks of a TF are considered as positive binding events. The negative set comprises *x* randomly generated, non-overlapping peaks, that have the same mean peak width as the positive peaks. The intersection between the positive and the negative set is the empty set. We compare the gold standard set to our TF predictions using bedtools intersect [53], with a minimum overlap of 1bp. All peaks in the negative set/positive set, that do not overlap any of our TF predictions are counted as True Negatives (TN)/False Negatives (FN). All predictions that do not overlap the positive set, are considered to be False Positives (FP). The overlapping predictions are evaluated with respect to TEPIC affinities and Fimo scores using the package pROC [22]. We report Precision-Recall AUC (PR-AUC) to measure method performance.

## 3 RESULTS

### 3.1 A segmentation-based method for gene expression prediction

In this work, we present a segmentation-based method to predict TF binding *in vivo*. The method can be applied to footprints as well as to open-chromatin and HM peaks. *TEPIC* has been tested on DNaseI-seq, and NOMe-seq data, although it is generally applicable to all open-chromatin methods, as long as open regions can be determined. Further, our method has been tested on Histone peaks for H3K4me3 and H3K27ac. In addition to the peak-centric view, the signal intensity of open-chromatin peaks is included in the TF binding prediction. We propose that incorporating the open-chromatin signal reflects the degree of openness of a particular genomic region in the cell pool of the considered sample. Hence, if certain regions are accessible in the majority of cells in a cell pool, higher weight is assigned to them by our method. In contrast to traditional hit-based methods, *TEPIC* is based on TF affinities to include low-affinity binding. We found that combining open-chromatin peaks, the signal intensity within those, and the consideration of low-affinity binding sites improve gene expression learning. Several aspects of our findings are detailed in the following sections.

### 3.2 Information about open-chromatin fraction in the cell population improves prediction

Recall from the Methods section that *TEPIC* has been tested with four different annotation setups to estimate TF affinities for genes: 3kb, 50kb, 3kb-S, and 50kb-S. Including the signal intensity within open-chromatin peaks improves the correlation between predicted and actual gene expression in both considered window sizes, as shown in Figure 2a. We observe that the performance of the different setups to summarise peak TF scores usually follows the order 3kb < 3kb-S < 50kb< 50kb-S. This holds except for the samples: K562, LiHe1, and LiHe2; there the 3kb-S setup performs better than the 50kb setup. However, combining exponential decay in the 50kb window and scaling with the open-chromatin signal outperforms all other tested variants. This might indicate that incorporating distal TF binding events is crucial to modelling gene regulation accurately. We also notice that scaling TF affinities with the open-chromatin signal seems to work better with DNaseI-seq (cell lines and primary human hepatocytes) than with NOMe-seq (T-cells). Additionally, note that the hepatocyte sample *LiHe2* performs worse than the other two hepatocyte replicates. This might be explained by the varying number of open-chromatin peaks between the replicates, as *LiHe2* is the replicate with the fewest open-chromatin peaks. In Supplementary Table 2, all learning results are shown.

**Figure 2.**
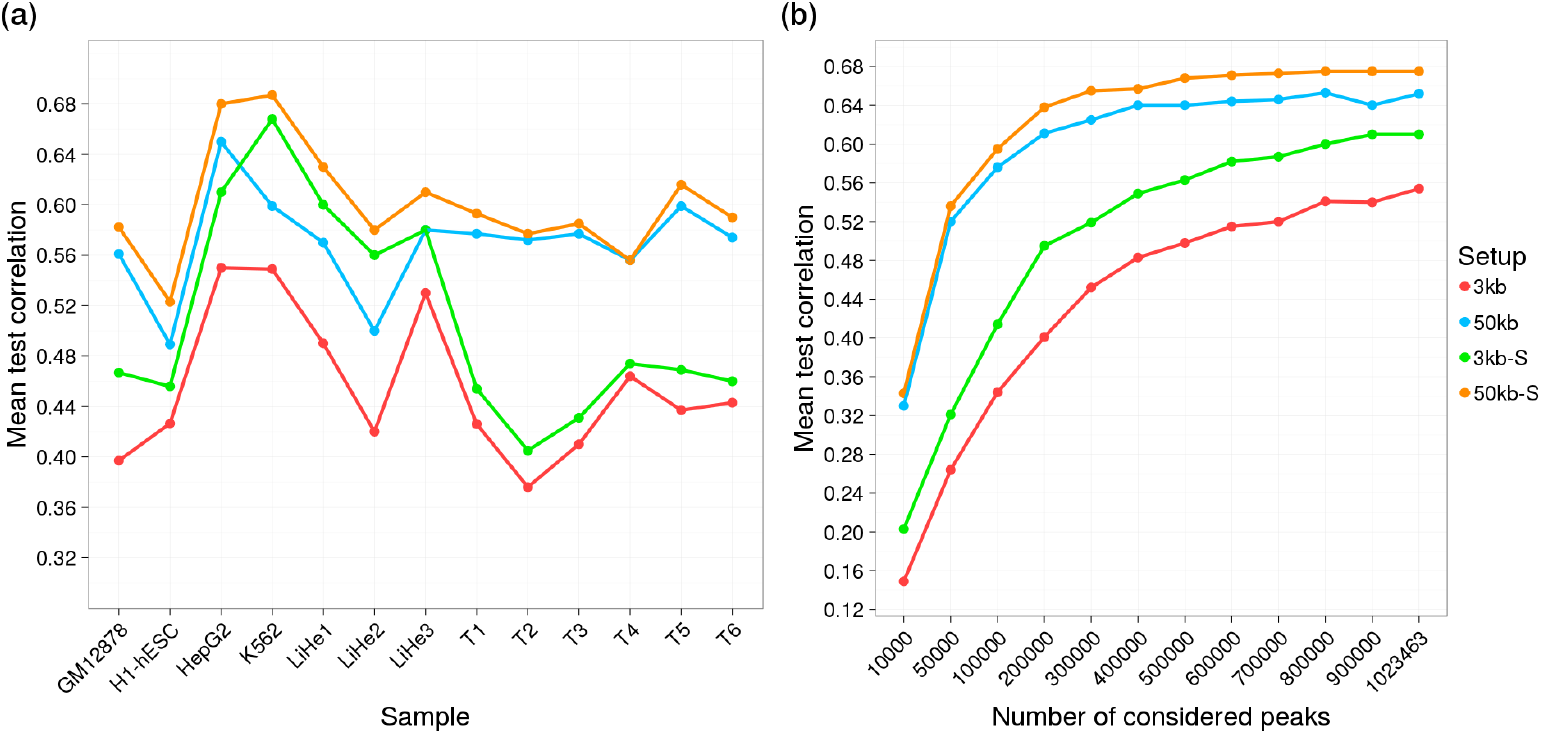
(a) Mean test correlation achieved in gene expression learning is shown for all tested setups and for all samples. The 50kb-S setup outperformed all other setups in all samples. We observe, that the scaling using the average peak intensity seems to work especially well for DNaseI-seq data, but not so well on NOMe-seq data, as the increase of the mean test correlation between 3kb and 3kb-S as well as between 50kb and 50kb-S is higher for the DNaseI-seq samples (GM12878, H1-hESC, HepG2, K562, LiHe1, LiHe2, and LiHe3) than for the NOMe-seq samples (others). (b) The learning performance for all setups with a varying number of considered peaks is shown. This analysis is based on HepG2 data only. An interesting observation is that the curves for the 50kb approaches saturate at around 400, 000 peaks, while the 3kb approach curves steadily increase till all peaks are included in the model.

### 3.3 Gene expression prediction depends on the number of open-chromatin peaks

We investigated the influence of the number of considered open-chromatin peaks on the performance of *TEPICs* prediction in the gene expression learning. For this purpose, twelve different peak sets using HepG2 DNase data were constructed according to the JAMM peak score. We considered 10, 000, 50, 000, 100, 000, 200, 000,…,900, 000, and all filtered peaks, 1, 023, 463. Interestingly, the performance of the 50kb and 50kb-S setups remains roughly constant for peak numbers ≥ 500000, while the performance of the 3kb and 3kb-S setups continuously increases until the end. This may be considered as support for the hypotheses that long-range regulation by TFs bound to distal binding sites is vital to modelling gene regulation. Additionally it can be seen that the difference between the setups pertaining to the same window size with and without the incorporation of the open-chromatin signal respectively, rises with increasing peak numbers. This might reflect the importance of prioritizing certain peak regions using the open-chromatin signal.

### 3.4 Including low-affinity binding sites improves over hit-based TF annotation

We compare hit-based TF scores with the affinity-based annotation used in *TEPIC*. As shown in Figure 4a, the incorporation of low-affinity binding sites using *TRAP* outperforms the traditional hit-based scores. Another advantage of *TEPIC* is that it consumes only 12.06 Gigabytes of memory, while the hit-based method *Fimo* required 86.6 Gigabytes (measured on HepG2).

**Figure 4.**
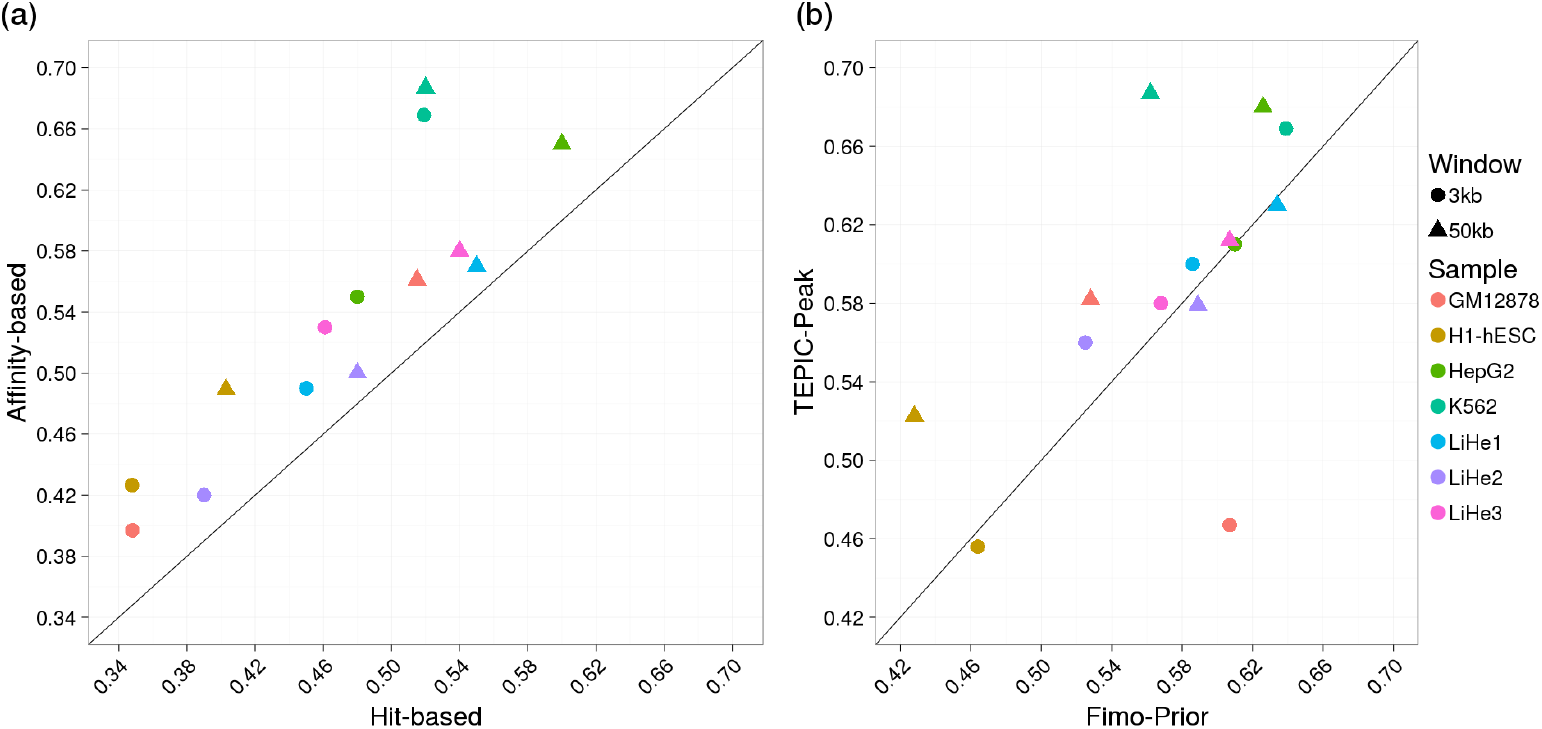
(a) The scatter plot shows the mean test correlation achieved in gene expression learning using TF affinity scores with TRAP and a hit-based peak annotation computed with Fimo. Clearly, the hit-based scores are outperformed by the TF affinities. (b) The scatter plot shows the mean test correlation achieved in gene expression learning using *TEPIC* applied on peaks and TF scores computed with Fimo-Prior. In general *TEPIC* scores show better performance in the expression prediction than those computed with *Fimo-Prior*, although both methods perform similar for several samples. Note that the scaled annotation versions of TEPIC are used in the comparison against Fimo-Prior.

### 3.5 Histone Marks contain information on TF binding

Histone Marks (HMs) have been successfully used in predicting TF binding sites [4, 13, 24, 51, 67]. Using ENCODE ChIP-seq data of H3K4me3 and H3K27ac obtained for HepG2, K562, GM12878, and H1-hESC we show that HMs can also be used in TEPIC. As shown in Figure 3, HMs lead to good performance in gene expression learning. Similar to the open-chromatin data, we note that using a larger window improves the learning results and that scaling the TF predictions using the abundance of the ChIP-seq peaks improves the results further in most cases, excluding the 50kb-S setup of H3K27ac in H1-hESC, HepG2, and K562. Further, we observe that H3K4me3 leads to better results than H3K27ac in all samples. This might be due to the strong association of H3K4me3 to active promoters [29], whereas H3K27ac is rather related to enhancer regions [11]. In particular, this might explain the reduced performance of H3K27ac peaks in the 3kb(-S) setups compared to H3K4me3. Additionally, we compared the performance of running *Fimo* in HM peaks to using TEPIC. Similar to the results shown in Figure 4a for DNaseI-seq data, we observed that the hit-based annotation is outperformed by TF affinities (Supplementary Figure 5).

**Figure 3.**
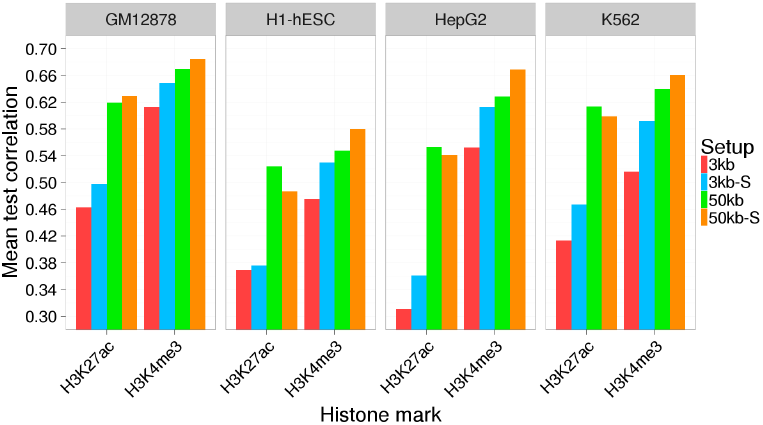
Gene expression learning results in GM12878, H1-hESC, HepG2, and K562 cells are shown for four different annotation setups using either the positions of H3K4me3 or H3K27ac peaks as input for TEPIC. Scores based on H3K4me3 work better than those based on H3K27ac across all samples.

### 3.6 TEPIC improves expression estimates compared to an epigenetic prior used with Fimo

We compare our approach against a state-of-the-art TF binding prediction method that extends *Fimo* with an epigenetic prior. We refer to this method as *Fimo-Prior*. It was shown to perform competitively to the earlier state-of-the-art *Centipede* [13].

We applied *Fimo-Prior* to DNaseI-seq data of HepG2, K562, GM12878, H1-hESC, LiHe1, LiHe2, and LiHe3. In Figure 4(b), we show the performance of *TEPIC* in the 3kb window (3kb-S) and the 50kb window (50kb-S) compared to *Fimo-Prior*. *Fimo-Prior* and *TEPIC* perform similar for both setups of LiHe1 and LiHe3, for the 3kb setup of HepG2, and for the 50kb setup of LiHe2. The 50kb setup of HepG2 as well as the 3kb setup of LiHe2 achieve better learning results, when *TEPIC* scores are used instead of *Fimo-Prior*. This also holds for all cell line samples excluding the 3kb setups of H1-hESC and GM12878.

In contrast to our observations presented in Figure 2, we observed that the performance of *Fimo-Prior* on K562, on H1-hESC, and on GM12878 decreased in the 50kb window compared to the 3kb window. For ChIP-seq data it was shown that extending the region up to 50kb improved the quality of gene expression prediction [49]. This effect might be due to the design of *Fimo-Prior*, which is a site-centric method that considers all binding sites in the 50kb window. Although the open-chromatin signal is used for reweighting, it may be that too many false positive hits are considered in the final gene TF scores. Overall, the performance of *TEPIC* is favourable compared to the performance of *Fimo-Prior*. We observed that the runtime of *Fimo-Prior* is extensive compared to *TEPIC*. Analysing the 50kb region for HepG2 using the prior of [13] took about 6.5 days, while *TEPIC* performs this task in 16 hours (using 16 cores), including the time required for peak calling with JAMM. We note however, that the current implementation of *Fimo-Prior* is not parallelized. A summary of runtimes recorded within this comparison is shown in Supplementary Table 5.

### 3.7 Footprints contain essential binding sites for gene expression prediction

So far, most *segmentation-based* methods identify TF binding sites by predicting footprints [24]. Here, we compared the footprint-based segmentation to a peak-based segmentation. To this end, we considered 452, 281 footprints in HepG2, 738, 707 footprints in K562, 598, 500 footprints in GM12878, and 1, 023, 559 footprints in H1-hESC identified with the currently most accurate footprinting method *HINTBC* [24]. We used TEPIC to annotate the regions around each footprint with a window of length 24bp and 50bp (see Methods). As the results between both setups are very similar, we present only the results for the slightly better 50bp setup and refer to Supplementary Figure 4 for a comparison of both. Figure 5 shows the comparison between *TEPIC* applied to footprints and peak regions. The peak-based approach outperforms the footprints in HepG2 and K562. In addition, peaks perform slightly better than footprints in H1-hESC. In GM12878, the footprint based approach outperforms the peaks.

**Figure 5.**
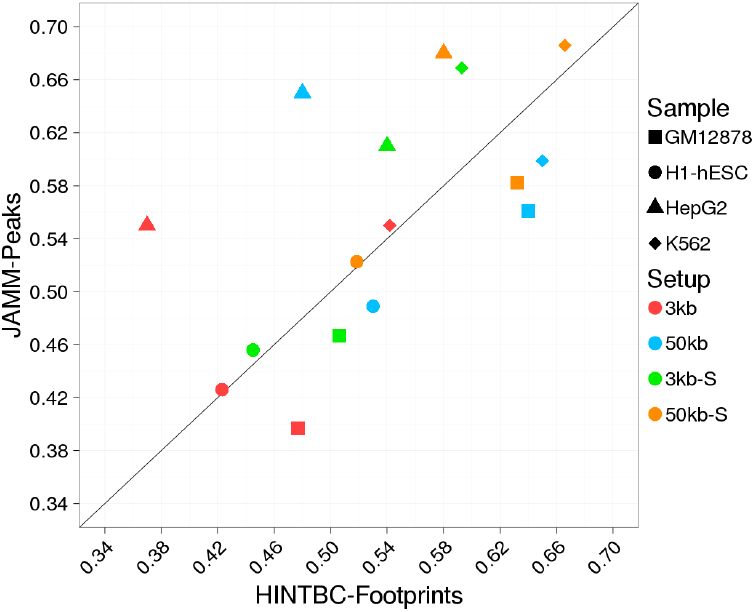
The scatter plot shows the mean test correlation achieved in gene expression learning using TF affinities computed within JAMM DNaseI-seq peaks and TF affinities computed within a 24bp window centred at footprints called using HINTBC. On HepG2 and K562, the peak-based approach outperforms the TF-footprints, whereas in GM12878 footprints lead to a better model performance On average, H1-hESC samples show a slighty better performance using peaks.

In addition, we see that incorporating the open-chromatin signal is also applicable to the extended footprint regions as the correlation increases between the 3kb and 3kb-S, as well as, between 50kb and 50kb-S approaches. Only for GM12878, the 50kb approach performs a little better than the 50kb-S approach. This observation also holds for the 24*bp* footprint extensions. Although using peaks to segment the genome seems to lead to better results on average, it is remarkable that the rather small footprint regions seem to cover most of the important binding sites. Using only 22.98%, 25.33%, 91.2%, and 36.02% of base pairs in footprinting regions compared to peak regions in HepG2, K562, GM12878, and H1-hESC respectively, illustrates that indeed most of the essential TF binding events in these cells are overlapping the footprint calls.

### 3.8 TEPIC applied to DNaseI-seq data performs comparable to TF ChIP-seq data in gene expression learning

We compared the performance of our method with that of gene expression learning using TF ChIP-seq data. In Figure 6 we show the learning results for HepG2, K562, GM12878, and H1-hESC. To illustrate the relation between the different TF binding prediction methods, the figure includes the best correlation achieved [1] on footprints, [2] using *Fimo* within open-chromatin peaks (labelled as Hit-based), and [3] using *Fimo-Prior*.

In HepG2 and K562, we find that *TEPIC* applied on peaks outperforms all other approaches, including *Fimo-Prior* as used in [46], and achieves correlation values that are close to what is obtained by using TF ChIP-seq data. In GM12878 and H1-hESC, TEPIC applied to footprints, outperforms the competitive approaches and also achieves good correlation. As the computational models lack some of the capabilities of the ChIP data, it was surprising to us that using a computational model, allows to get so close to ChIP-seq based predictions for some of the datasets. In addition to comparing all pwms against all available ChIP-seq data, we compared the performance of using exactly those pwms for which ChIP-seq data is available and vice versa. Although the overall correlation between observed and predicted gene expression decreased, we again found that *TEPIC* produces results often close to those with ChIP-seq data (Supplementary Figure 6).

**Figure 6.**
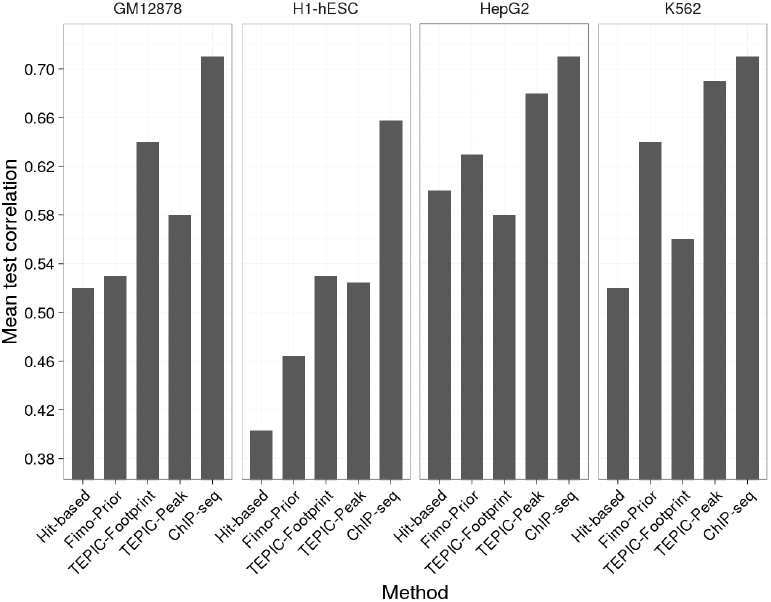
Barplots showing the performance of gene expression learning for HepG2, K562, GM12878, and H1-hESC using several different computational TF scores as well as TF-ChIP-seq data. Although the ChIP-seq data outperformed all computational TF binding prediction methods, *TEPIC* scores achieved good results compared to all other computationally derived scores. In this Figure, the best performing variants of the individual methods are represented.

### 3.9 TF binding predictions computed by TEPIC perform well in a comparison to TF-ChIP-seq data

The common way to evaluate TF binding prediction methods is to conduct a comparison to TF ChIP-seq data. We used such an evaluation setup to benchmark the different approaches in addition to the analysis of gene expression prediction. To this end, we calculated Precision-Recall(PR) AUCs, as described in the Methods section, for predictions on HepG2, K562, GM12878, and H1-hESCs compared to TF ChIP-seq data. We compared *TEPIC* applied to open-chromatin peaks against *Fimo* scores computed in open-chromatin peaks, against *Fimo-Prior*, which is applied genome-wide, and against *TEPIC* scores computed in footprints. Detailed results are shown in Supplementary Figures 8, 9, 10, and 11. In Table 1 we present our results in a compact way, by listing the mean PR-AUC values over all TFs for the individual comparisons.

**Table 1.**
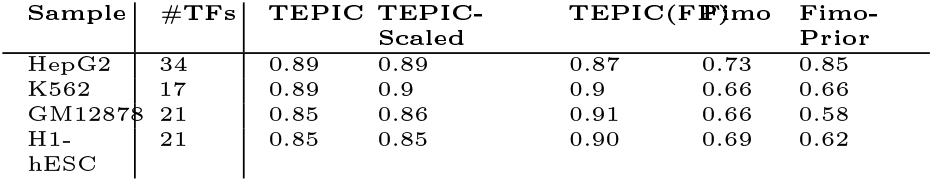
Mean Precision-Recall(PR)-AUC of computational TF predictions compared to experimentally determined TF binding sites. We consider *TEPIC* applied to peaks and footprints(fp), a scaled and an unscaled version of *TEPIC*, *Fimo* applied to peaks, and *Fimo-Prior*.

We observe that the scaled *TEPIC* scores perform comparable to the unscaled scores, except for a minor improvement in K562 and HepG2. This indicates that prioritising peaks using the open-chromatin signal is more relevant in a gene expression prediction task compared to an evaluation against TF ChIP-seq data. The ChIP-seq comparison clearly indicates, that affinity-based scores are superior to a simpler hit-based annotation using *Fimo*, as mean PR-AUC values across samples are considerably larger for *TEPIC* scores than for *Fimo*. This is in concordance with the analysis shown in Figure 4a. The mean PR-AUC values computed for Fimo-Prior are superior to the simple Fimo scores only on HepG2, in K562 they are equal and worse for the remaining samples. TEPIC scores computed in footprints and peaks show a comparable performance, which is in concordance to the findings shown in Figure 5.

### 3.10 Models learned using TEPIC scores are tissue-specific

To determine whether the learned models are tissue-specific, a Principal Component Analysis (PCA) was performed on the model coefficients of all samples used in this study. As it can be seen in Figure 7, the primary human hepatocyte samples (LiHe) are clearly separated from the remaining samples, while HepG2, a human liver cancer cell line, is their next neighbour, according to PC1. The T-cell samples are positioned in the right area of the PCA plot. Their nearest neighbour is GM12878 which is located very close to two of the T-cell samples. GM12878 is a lymphoblastoid cell line.

**Figure 7.**
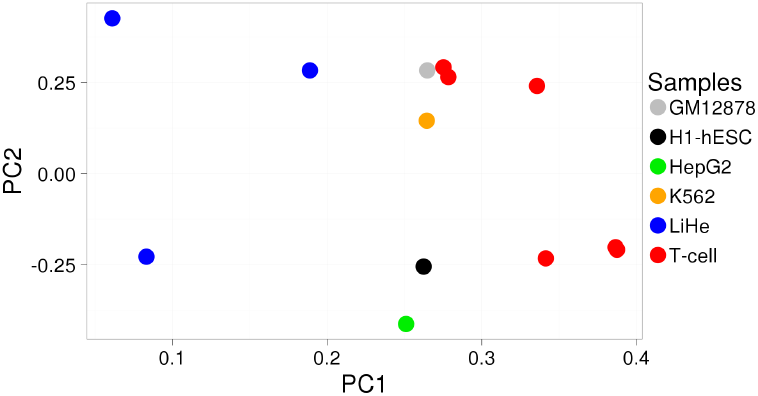
Principal component analysis of normalised model coefficients for all samples considered in this study. There is a clear separation of primary human hepatocytes, cell lines, and T-cells.

Lymphoblasts can differentiate into T-cells, hence the position of GM12878 in the PCA plot could be explained. We note however, that the T-cell samples are obtained from NOMe peaks, whereas all other peaks are from DNAse1, therefore PC1 appears to also capture that difference.

In addition to the PCA analysis, we performed a cross-sample comparison using our models. To this end, we learned a model using data for a distinct sample *x* and used this model to predict gene expression across all samples. The results are shown as a heatmap in Supplementary Figure 7. Similar to the PCA analysis, this experiment argues for a tissue-specificity of our models, as the clustering of the model performances clearly indicates a similarity/dissimilarity between related/unrelated cell types. Thus, it should be worthwhile to investigate the feature vectors in more detail to learn about tissue-specific regulators.

### 3.11 TF expression filtering does not reduce model performance and simplifies interpretation

We checked how many of the TFs selected as a non-zero feature by the elastic net model are actually being expressed. Thereby, we found that the mean expression level of selected TFs is higher than the mean expression level of the TFs that are not selected (Supplementary Figures 12 and 13). Therefore, we repeated the gene expression learning with a set of TFs that has been filtered with regard to expression levels. We used a low FPKM cut-off of 1.0 and additionally removed all TFs that could not be mapped to a gene ID. Supplementary Figure 14 shows that this reduction of considered TFs does not reduce the learning performance. As the TF filtering reduces the number of features, it simplifies the interpretation of the model coefficients. Non-zero coefficients mean that TFs influence gene expression, either as activators (positive coefficients) or as repressors (negative coefficients). *TEPICs* different annotation setups allow us not only to estimate the influence of different TFs on gene expression but also to compare factors that are predicted to bind in the promoter region (3kb-S setup) and those that are predicted to bind in addition to distal regions to the TSSs of genes (50kb-S setup). Thus, we will consider both setups in the analysis of the primary human hepatocytes and the T-cell samples described in the following sections.

### 3.12 Analysis of primary human hepatocyte datasets using DNaseI-seq data

To investigate the role of TFs in the liver hepatocyte samples, we computed the total feature overlap between the learned models. In Figure 8a a Venn diagram is shown visualising the overlap between the models. We found that 65 (38.5%) TFs are commonly selected between all replicates using the 50kb-s setup (Figure 8a). In Figure 8b, we show the top 10 positive and top 10 negative features selected by our model. By conducting literature research we found that there is evidence for 52 of the 65 factors to be associated to hepatocyte function. Within the top 10 positive and negative features, we found for example, the heterodimer *PPARG::RXRA*. This factor plays a key role in hepatic transcription [52]. Another example is *CEBPA*, which is known to be important in liver regeneration [10, 14]. The TF *GATA4* was shown to be involved in liver induction [2]. *CTCF* was found to have a role in imprinting liver [23, 28], and *NRF1* has a protective function against oxidative stress in liver [68].

**Figure 8.**
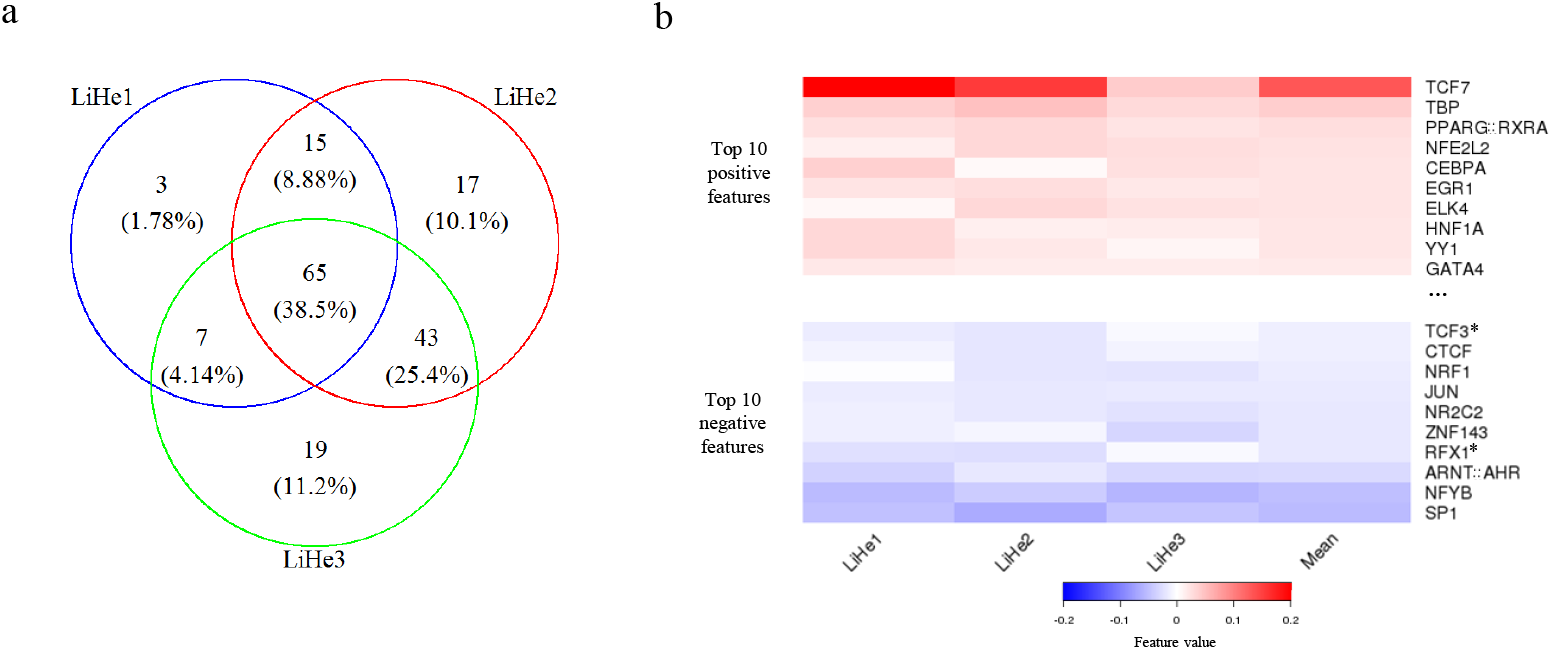
(a) Venn diagram visualising the overlap between the liver hepatocyte replicates using the 50kb-S annotation. In total, 65 factors are shared between the replicates, and only 3, 17, and 19 are selected uniquely. (b) Heatmap listing the top 10 positive and top 10 negative selected features, which are among the 65 shared features in the 50kb-S setup. TFs labelled with a * could not be validated by literature to be related to hepatocytes.

A list of all factors is provided in Supplementary Table 3, Supplementary Figure 15 is analogous to Figure 8 but based on the 3kb-S annotation.

### 3.13 Application to NOMe analysis in T-cells

Overall, there are 53 (39%) TFs commonly selected in all T-cell samples. The feature overlap between the individual T-cell replicates is shown in Figure 9a. We suggest that those 53 TFs are potential key regulators within T-cells. By conducting literature research, we found evidence that 42 out of the 53 are known to be related to the immune system, see Supplementary Table 5. For example, among the top 10 positive and negative coefficients (Figure 9b) we found the factor *Gmeb1*. This factor was shown to inhibit T-cell apoptosis [33]. Another TF with a positive coefficient is *Ets1*, which was shown to be critical for T-cell development [17]. Among the negative coefficients is the factor *Zbtb7b*, which is known to act as a repressor in CD4+ T-cells [65].

**Figure 9.**
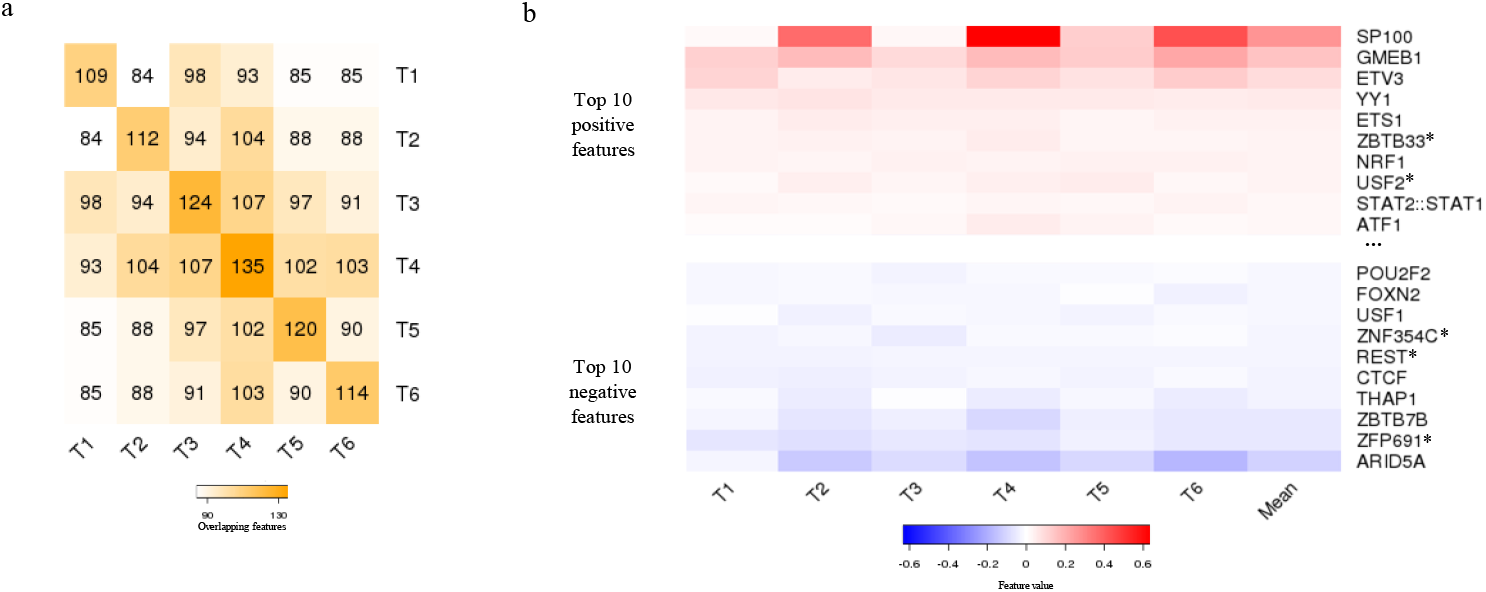
(a) Heatmap showing the overlap between the T-cell replicates. There are 53 (39%) factors shared between all T-cell samples. (b) The top 10 positive and top 10 negative features among the 53 shared ones, are listed here. TFs labelled with a * could not be validated by literature to be related to regulation in T-cells. For the others, we were able to find literature that sets those factors into relation to T-cells (see Supplementary Table 4).

By comparing the TFs selected between the 3kb-S and the 50kb-S setup (see Supplementary Table 4) we observed that the TF *TBP*, which binds to the TATA motif in core promoters, is selected only in the 3kb-S setup. This might indicate that factors, that are involved in basal transcriptional regulation, such as *TBP* [37] might not contribute additional information to the model if distal binding events are considered. We also noted that the feature coefficient signs agree between all TFs common in both setups. This can be seen as a hint to the robustness of the learning itself. Supplementary Figure 16 shows these analysis for the 3kb-S setup on the T-cells.

## 4 DISCUSSION

Here, we introduce a new method, *TEPIC*, to predict TF binding using an open-chromatin assay as a prior to reduce genomic search space. Within *TEPIC*, several new aspects in this field are proposed.

Previous segmentation-based methods for TF prediction segment the genome using TF footprints [24]. Here, we include a segmentation paradigm which we call *peak-centric*, as we consider all open-chromatin peaks to represent accessible DNA and predict TF binding exactly in these regions. Earlier, it was observed that DNaseI-seq signal corresponds well to TF-binding, e.g. in [13], but a peak-centric segmentation has not been explored in detail, so far. A comparison to footprints called with *HINTBC* in a gene expression learning setup showed that peaks perform similar to footprints. A clear advantage of the peak-centric paradigm is that it is assay-independent. We applied our method to DNaseI-seq data, which is the open-chromatin assay used in the majority of TF binding prediction methods, but also to NOMe-seq data, without any changes to the code. This is not easily possible for footprint-based methods, as they are assay-specific. *TEPIC* could be easily applied to other open-chromatin assays, for example ATAC-seq data [6].

An investigation whether the performance of a peak-centric segmentation would be affected by the used peak caller showed that *JAMM* [31] peaks deliver better results on DNaseI-seq data than MACS2 [70] peaks. However this could have been expected, as *JAMM* was designed to handle the characteristics of DNaseI-seq data. The learning results with MACS2 peaks are listed in Supplementary Table 2 and are visualised in Supplementary Figure 3.

In order to improve TF binding predictions further, we included the absolute signal of the open-chromatin assay within a peak in the score describing TF binding (see Methods). This allows us to capture heterogeneity of TF binding over the large amount of cells considered in bulk sequencing approaches. We showed that this extension improves gene expression prediction compared to the 3kb and 50kb approaches (Figure 2a). Therefore the biological interpretation of the models becomes more reliable. The scaling also improved predictions carried out on footprints. In addition to considering the open-chromatin based segmentation, we have shown that also HMs can be used within *TEPIC* to identify candidate TF binding sites and that incorporating the signal within the HM-peaks also improves gene-expression prediction.

Former TF binding prediction methods that integrate open-chromatin information were using a hit-based approach and had to rely on p-value thresholds. It is not obvious that estimating binding affinity of a TF, *e.g.* using *TRAP* [55], within the complete peak regions must improve over a more reduced search space when using hit-based methods to define binding sites within peaks, as one could argue that the affinity based approaches accumulate more noise. We believe that there are two major reasons why *TRAP* outperforms the hit-based approach: first by default the same p-value threshold is used for all pwms, although the information content of pwms may vary widely. An additional optimization of the p-value threshold for each pwm may improve the result. Second, a drawback of hit-based methods is that low-affinity binding sites are lost. Incorporating these biologically important binding events [12, 59] seems to be relevant for improving the predictions.

The combination of those novel aspects enabled *TEPIC* to outperform a state of the art site-centric method that incorporates an epigenetic prior within *Fimo* [13]. *TEPIC* achieves the best correlation in gene expression learning among all tested methods and nearly reached the quality of using several ChIP-seq data sets. However, these findings also point us to a few drawbacks of our method. Although the exponential decay in the 50kb window, proposed in [49], improves the learning result, it is likely that it also adds noise to the gene TF scores. This could be improved by replacing the exponential 50kb weighting with a more sophisticated function based on 3D chromatin structure using Hi-C data. In addition, the pwm based annotation allows neither modelling indirect TF binding nor allows a modelling of TF complexes. These points might explain why we cannot fully reach, or even overcome, the quality of ChIP-seq based predictions.

*TEPIC* is an unsupervised method for predicting TF binding. Because we wanted to include as many TFs as possible in the input for the gene expression learning, we decided to exclude supervised methods, such as the recently published *BinDNase* [32], in the comparison with other methods. These approaches require the presence of ChIP-seq data for all TFs of interest and therefore are not applicable for many of the large epigenetic datasets produced.

To test the performance of *TEPIC’s* TF predictions, we performed an evaluation against TF-ChIP-seq data as well as gene expression prediction experiments. Note, that we do not conduct a TF motif filtering to remove ChIP-seq peaks that are unlikely direct binding events of the chipped TF. For this step, either Fimo or TEPIC predictions would normally be used as a filtering criteria, leading to a bias in the evaluation setup. The rather bad performance of Fimo-Prior in our TF-ChIP-seq evaluation might be due to the design of our gold standard set in a peak-centric manner or because the prior is not well suited to be applied genome-wide. For example, the number of stored hits is limited in the current implementation of Fimo, which might cause problems if the tool is used on a large scale. However, it was shown by both evaluation strategies that a hit-based TF annotation is less accurate than an affinity-based annotation. Moreover, both validation setups encourage a deeper analysis comparing TF annotations based on either open-chromatin peaks or footprints, as it is not obvious which segmentation methodology is more accurate in general. As footprint calling is computationally more involved than peak calling, the latter might be more applicable in practice. As pointed out in [58], TFs with short DNA residence times do not exhibit footprints, therefore it might be possible to improve predictions by deciding for each TF whether peaks or footprints should be used.

We note that gene expression learning for validation has several advantages over a simple comparison to ChIP-seq data. As it was observed by the authors of *Millipede* [42], there is no common strategy of validating TF binding predictions directly by comparing them to TF ChIP data. Using gene expression learning [1] avoids problems arising by imbalanced positive and negative sets, and [2] vague definitions of gold standard sets, and [3] enables a biological interpretation of the results. We believe the method may be exploited in other aspects relevant for TF binding prediction, *e.g.*, the evaluation of footprinting methods [24].

In this study, we applied our method to primary cell types, primary human hepatocytes and CD4+ T-cells, as well as to cell lines. We showed that the TF binding predictions of *TEPIC* used for gene expression learning led to the identification of TFs that are highly associated with the regulation of the analysed cell types and identified a number of interesting candidates that show strong regulation but are not associated with regulation in these cells.

The observation that factors which are generally participating in transcriptional regulation at promoters, such as TBP [37], are not stably selected by the learning method applied to the 50kb window, suggests that these are not more predictive for gene expression than factors that bind in more distal regions from TSSs, *e.g.*, enhancer regions that are known to define tissue-specificity.

## 5 CONCLUSION

We propose a novel method for TF binding predictions, validated using gene expression learning. Compared to previous segmentation-based methods, our method offers a peak-centric mode and, thus, is assay-independent. Instead of using a hit-based annotation, *TEPIC* uses an affinity-based annotation, and additionally combines TF affinities with the open-chromatin signal in a simple quantitative manner to improve the binding predictions further. We showed that with just a single open-chromatin assay and straightforward data preprocessing, it is possible to achieve approximately the same quality in gene expression learning as compared to the use of several expensive ChIP-seq assays. Further *TEPIC* outperforms several competitive approaches. Our method including routines for parallelization is freely available at *www.github.de/schulzlab/TEPIC*.

## Acknowledgments

We thank all labs that contributed to the ENCODE data used in the scope of this project. We thank Eduardo G Gusmao and Ivan G Costa for providing us with the footprint calls in HepG2, K562, GM12878, and H1-hESC using their method *HINTBC*. We thank Patricio Godoy for help with literature search for liver TF regulation. We thank Matthias Heinig for creating the R-package of TRAP.

